# *Candida albicans* enhance *Staphylococcus aureus* virulence by progressive generation of new phenotypes

**DOI:** 10.1101/2024.06.26.600854

**Authors:** Betsy Verónica Arévalo-Jaimes, Eduard Torrents

## Abstract

*Candida albicans* and *Staphylococcus aureus* have been co-isolated from several biofilm-associated diseases, including those related to medical devices. This association confers advantages to both microorganisms, resulting in detrimental effects on the host. To elucidate this phenomenon, the present study investigated colony changes derived from non-physical interactions between *C. albicans* and *S. aureus*. We performed proximity assays by confronting colonies of the yeast and the bacteria on agar plates at six different distances for 9 days. We found that colony variants of *S. aureus* originated progressively after prolonged exposure to *C. albicans* proximity, specifically in response to pH neutralization of the media by the fungi. The new phenotypes of *S. aureus* were more virulent in a *Galleria mellonella* larvae model compared to colonies grown without *C. albicans* influence. This event was associated with an upregulation of *RNAIII* and *AgrA* expression, suggesting a role for α-toxin. Our findings indicate that *C. albicans* enhances *S. aureus* virulence by inducing the formation of more aggressive colonies.

**Importance:** For decades, it has been known that *C. albicans* increase *S. aureus* virulence, resulting in a “lethal synergism”. However, it was only recently identified that this outcome is driven by the sustained activation of the staphylococcal *agr* system in response to *C. albicans* environmental modifications. Our experimental design allowed us to observe individual changes over time caused by the proximity of both microorganisms. As a result, we report for first time that *C. albicans* exposure induces the generation and favors the growth of *S. aureus* colony variants with increased expression of virulence factors. Our findings highlight the importance to understanding the intricate connection between environmental responses, virulence and fitness in *S. aureus* pathogenesis.

## Introduction

In the past, Koch’s postulates helped identify the etiological causes of several monomicrobial infections (1). Then, high-throughput genome sequencing techniques significantly expanded the notion of microbial biodiversity harbored within the body, revealing, for instance, more than 1000 species residing in the gut (1). Currently, it is recognized that the coexistence of multiple microorganisms in the same host niche fosters interactions among them that can drive disease onset and outcome (1). Furthermore, these complex communities can adopt the biofilm mode of life, secreting an extracellular matrix that surrounds and protects them from environmental attacks, including antimicrobial agents (1, 2)

Among the polymicrobial infections, those involving the presence of fungi and bacteria are particularly significant in the clinical context (2). The yeast *Candida albicans* and the Gram-positive bacterium *Staphylococcus aureus* are among the most prevalent nosocomial pathogens responsible for severe morbidity and mortality (3). This duo is frequently present in intra-abdominal infections that can disseminate and produce systemic infections that are very difficult to treat (1, 2). Moreover, *C. albicans* and *S. aureus* have been co-isolated from several biofilm-associated diseases, including chronic wound infections, cystic fibrosis, urinary tract infections, and medical-device-related infections (1, 2). Being part of these polymicrobial biofilms confers several advantages to both members, such as enhanced tolerance to vancomycin in *S. aureus* (4) and to miconazole in *C. albicans* (5). Likewise, the addition of conditioned medium from *S. aureus* dramatically increases *C. albicans* biofilm growth (6).

The cooperative relationship of these microorganisms involves both physical interactions and interspecies communication (2, 7). The chemical interactions between *C. albicans* and *S. aureus* include secreted molecules from the quorum sensing (QS) system (1, 2). In *S. aureus*, the best-characterized QS system is the accessory regulatory (*agr*) system, consisting of the promoter P2 and P3 that drive the expression of the transcripts ARNII and ARNIII, respectively (1). RNAII is encoded for four genes *agrA*, *agrB*, *agrC* and *agrD*. AgrB is a membrane protein that modifies and secretes the pre-signal peptide AgrD into its mature form, the autoinducing peptide 2 (AIP-2). AIP-2 is recognized by the membrane-bound protein kinase AgrC, which phosphorylates AgrA, activating P2 and P3. RNAIII is the effector molecule of the *agr* system and is responsible for toxin production, including α-toxin, a fundamental tool in *S. aureus*’ pathogenicity (1).

Polymicrobial infections are often associated with poor patient prognosis (7, 8). In the case of *C. albicans* and *S. aureus*, it has long been known that their intraperitoneal coinfection has a synergistic effect on mouse mortality (9). Increased virulence of *C. albicans* and *S. aureus* co-infection has also been reported in zebrafish embryos, *Galleria mellonella* larvae, and patients with systemic infections (10, 11). However, it was only recently that the major lethality driver and its mechanism of activation were identified. *C. albicans* ribose catabolism and alkalinization of the extracellular pH led to sustained activation of the staphylococcal *agr* system and the production of the effector molecule α-toxin (3, 12, 13). α-toxin can cause membrane damage, lysis, eicosanoid stimulation, disruption of tight junctions, activation of platelet aggregation, and dysregulation of the hemostatic system, leading to organ failure (1).

Further studies elucidating the details of the "lethal synergism" among this interkingdom association will aid in the development of new therapeutic targets. As the understanding of microbial interactions makes it clearer that the successful treatment of polymicrobial infections will arise from targeting the community as a whole (6). For this reason, the present study aimed to investigate colony changes resulting from non-physical but close interactions in a long-term *C. albicans* and *S. aureus* co-culture. To that end, proximity assays confronting the yeast and bacteria at six different distances for 9 days were developed. We found that *S. aureus* virulence enhancement results from progressive phenotypical diversification of the original colonies.

## Results

### *C. albicans* proximity promotes the generation of *S. aureus* colony variants

We observed non-physical interactions between the well-known opportunistic pathogens *S. aureus* (Sa) and *C. albicans* (Ca) through a proximity assay spanning six different distances over 9 days (Fig. 1). Inoculum drops deposited on the agar allowed for the development of well-defined colonies of both microorganisms, which were visible from day 1. In the case of *C. albicans*, colonies expanded progressively over the days. However, in the control (Ca – Ca), the size of the colonies decreased with increasing proximity, while in the proximity assay (Ca – Sa), the size remained constant across positions (Fig. S1).

**FIG 1.**
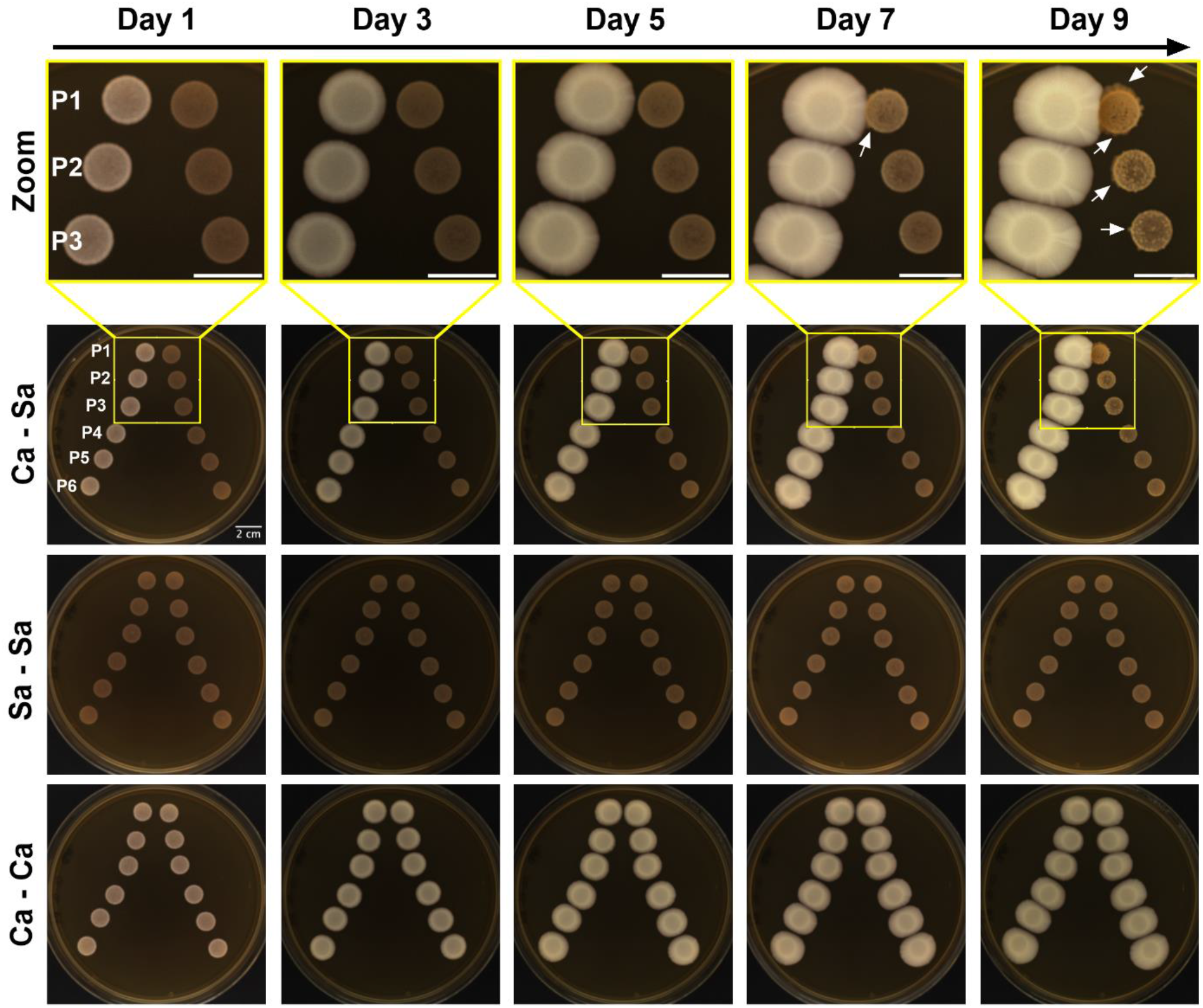
Proximity assay timeline of *C. albicans* (Ca) and *S. aureus* (Sa). Ca – Sa depicts zoomed (top) and original images (bottom) of *C. albicans* colonies (left) developed at six different distances (P1= 1 cm, P2= 1.5 cm, P3= 2 cm, P4= 3 cm, P5= 4 cm and P6= 5 cm) from *S. aureus* colonies (right) monitored over 9 days with a 1-day interval. White arrows indicate evident growth of new phenotypes within *S. aureus* original colonies. Sa – Sa represents the control condition in which *S. aureus* inoculum was used in both columns, while Ca – Ca is the control condition where *C. albicans* inoculum was used in both columns. The displayed images are representative of three different experiments and were processed with ImageJ software. The scale bar of 2 cm applies to all cases.

In contrast, the size of *S. aureus* colonies in the proximity assay (Ca – Sa) remained constant (Fig. 1). However, between day 5 and 7, small points of additional growth of *S. aureus* were observed within the original colony (see Fig. 1 small white arrows). The quantity and size of the growth spots increased over time, leading to fusion, and covering the original colony entirely. By days 7-9, the expansion of new *S. aureus* colonies continued and extended beyond the edges of the original colonies (see Fig. 1 small white arrows). Interestingly, the emergence of new phenotypes was linked to the proximity to *C. albicans*, where the process of generating new variants was faster at the closest position (P1= 1 cm). Consequently, the effect gradually decreased over subsequent positions, being minimal in the last ones (P5= 4 cm and P6= 5 cm) and absent in the control plate (Sa – Sa).

The described production of new *S. aureus* colonies was absent in proximity assays with other bacteria but were present with *Candida parapsilosis* (Fig. 2), indicating that the colony generation effect could be a specific interaction of *S. aureus* with the genus *Candida*.

**FIG 2.**
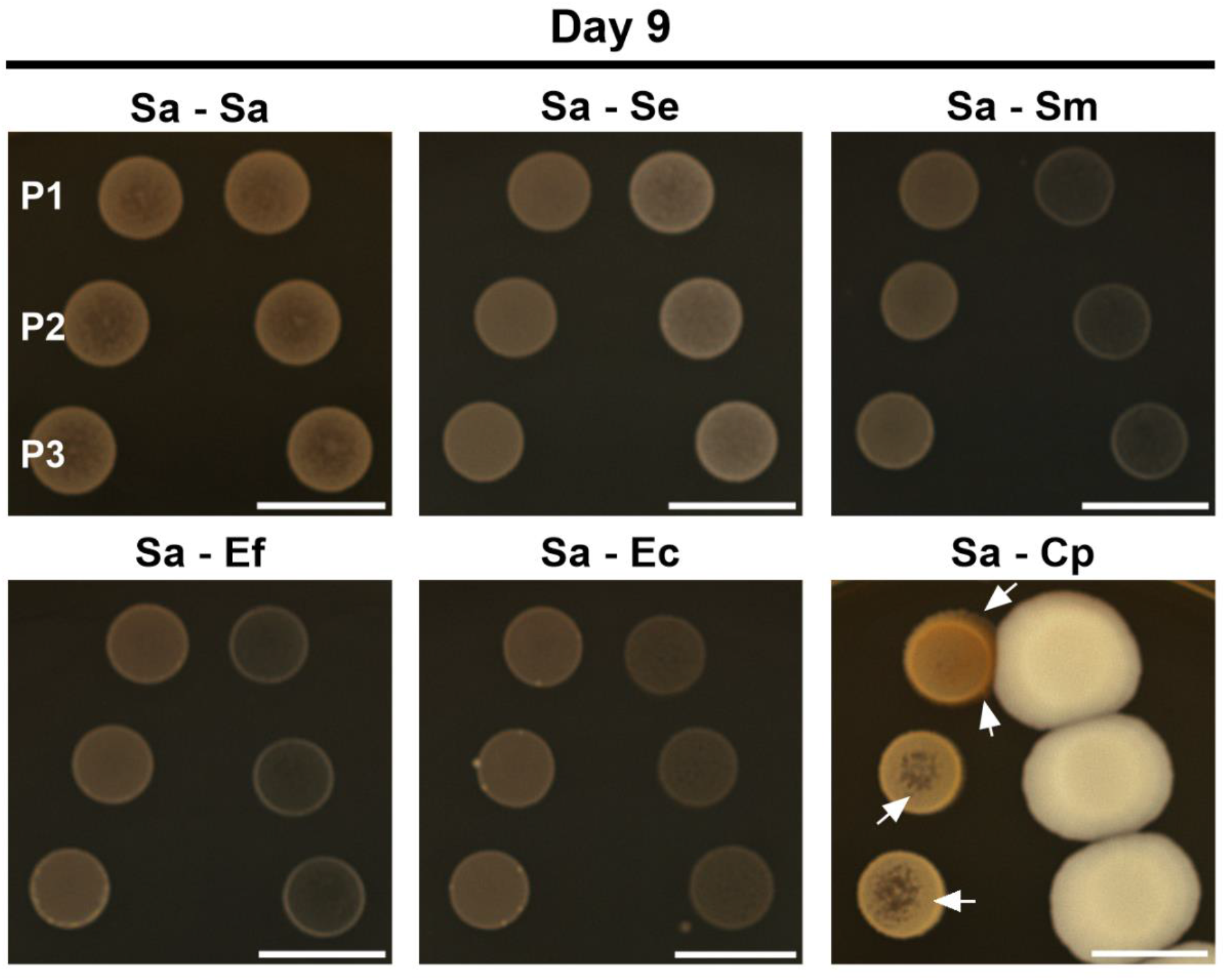
Proximity assay of *S. aureus* (Sa) with other microorganisms. Zoomed images of *S. aureus* colonies (left) with itself (Sa – Sa), *S. epidermidis* (Sa – Se), *S. mutans* (Sa – Sm), *E. faecalis* (Sa – Ef), *E. coli* (Sa – Ec) and *C. parapsilosis* (Sa – Cp) (right) at day 9. Displayed images are representative of three different experiments and were processed with ImageJ software. The scale bar corresponds to 2 cm and applies to all cases. The shown distances correspond to P1= 1 cm, P2= 1.5 cm and P3= 2 cm.

### Generation of *S. aureus* colony variants depends on pH and initial glucose levels

Considering the ability of *C. albicans* to alkalinize the media and the influence of pH on *S. aureus* virulence (13–16), we evaluated whether the appearance of *S. aureus* colony variants induced by *C. albicans* were related to pH changes. To this end, we performed the proximity assay by adding the pH indicator Bromocresol purple to the YPD agar medium (Fig. 3). The Sa – Sa control revealed that *S. aureus* growth from day 1 was accompanied by a pH decrease, which persisted throughout the experiment. In contrast, Ca – Ca control demonstrated that *C. albicans* growth led to a progressive increase in pH. These individual behaviors were maintained when both microorganisms were grown together (Ca – Sa). However, we observed that after day 5, *C. albicans* proximity to *S. aureus* nullified bacterial acidification. This alkalinization resulted in a predominance of a neutral pH in the positions with closer proximity between the species (P1-P3) by day 9. Though, the most meaningful finding, was the perfect overlap between *C. albicans* pH modulation and the origination of *S. aureus* colony variants (Fig. 3). Thus, we tested if *C. albicans* induced the formation of colony variants in other bacterial species, finding positive results with *S. epidermidis* and *E. coli* (Fig. S2).

**FIG 3.**
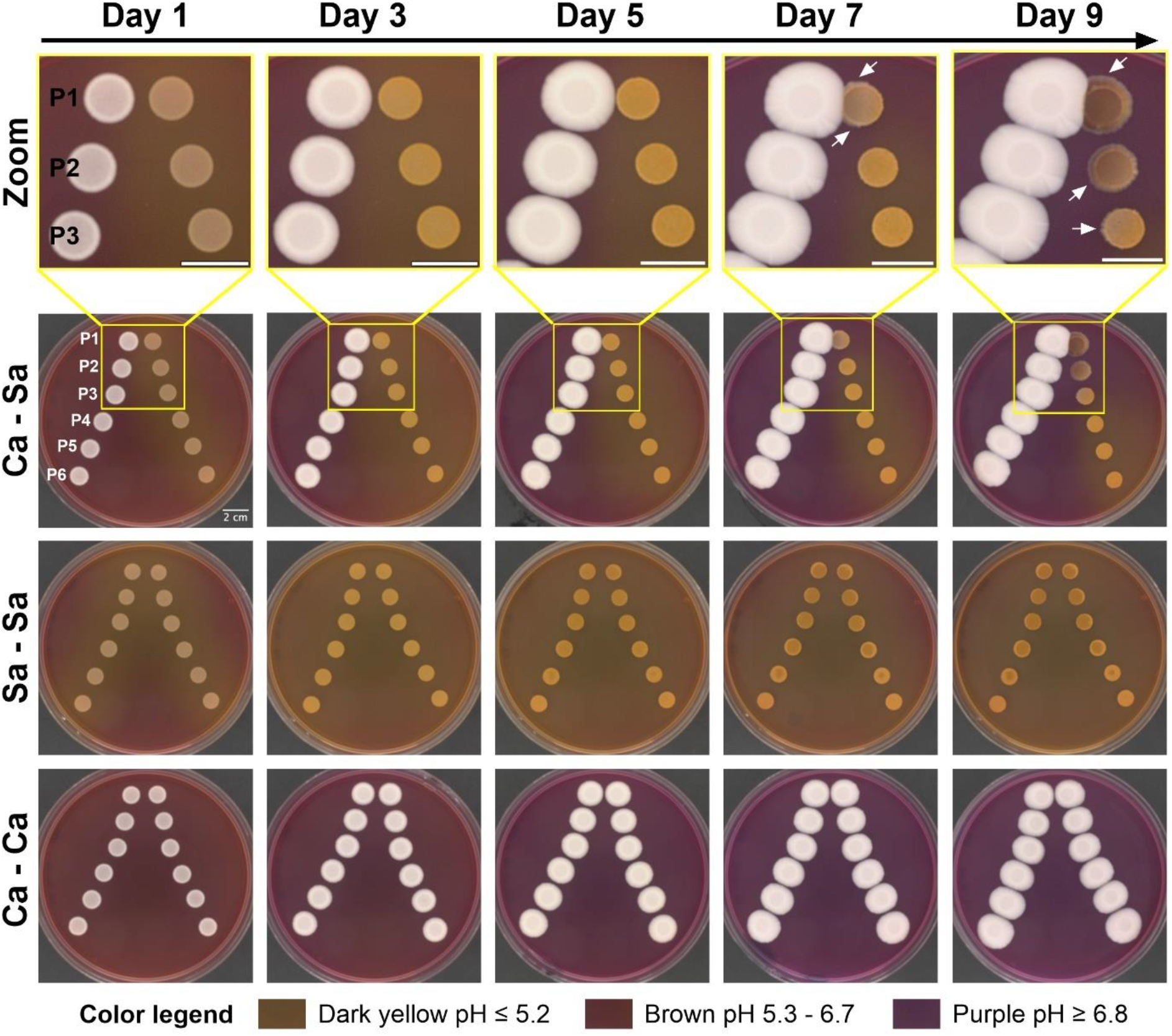
Timeline of pH changes in a proximity assay of *C. albicans* (Ca) and *S. aureus* (Sa). Ca – Sa depicts zoomed (top) and original images (bottom) of *C. albicans* colonies (left) developed at six different distances from *S. aureus* colonies (right) in YPD agar with bromocresol purple. Plates were monitored over 9 days with a 1-day interval. White arrows indicate evident growth of new phenotypes within *S. aureus* original colonies. Sa – Sa represents the control condition in which *S. aureus* inoculum was used in both columns, while Ca – Ca is the control condition where *C. albicans* inoculum was used in both columns. The displayed images are representative of three different experiments and were processed with ImageJ software. The scale bar of 2 cm applies to all cases. The shown distances correspond to P1= 1 cm, P2= 1.5 cm, P3= 2 cm, P4= 3 cm, P5= 4 cm, and P6= 5 cm.

Next, we assessed if the sole action of pH alkalinization was sufficient to induce the generation of *S. aureus* colony variants. Thus, we exposed *S. aureus* colonies grown in a proximity assay configuration to increasing quantities of NaOH 2M for 7 days (Fig. 4A). Again, we observed the appearance of *S. aureus* colony variants in positions (P1-P3) with greater exposure to pH changes. As expected, the PBS control did not induce either pH changes or generation of *S. aureus* variant colonies. This finding showed that the presence of *C. albicans* is not necessary to obtain the phenotypic variants of *S. aureus* even though in the case of proximity assays is the responsible of this pH change.

**FIG 4.**
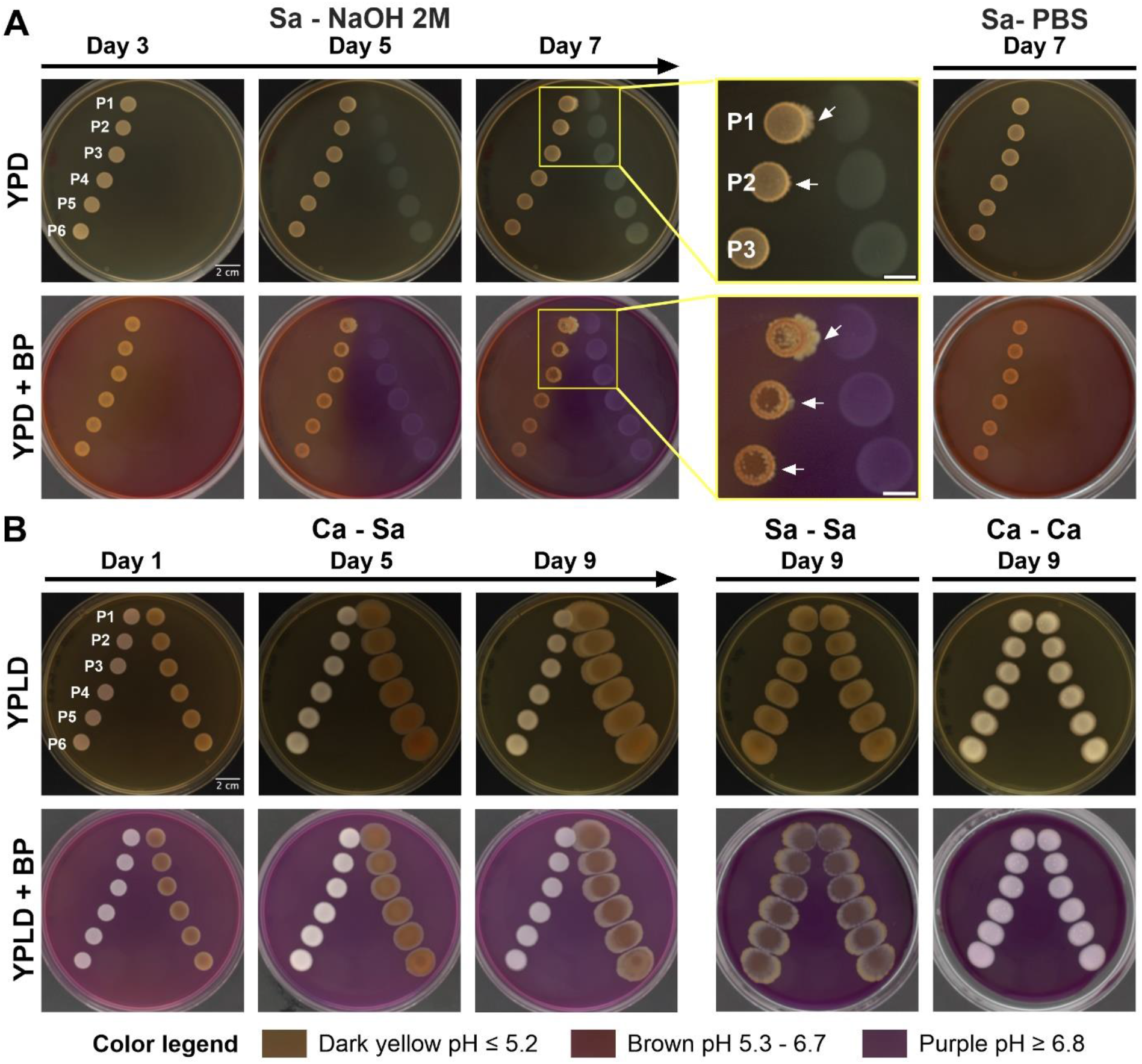
pH and glucose dependence in the generation of *S. aureus* colony variants. (A) Timeline of the effect of *S. aureus* colonies’ exposure to NaOH 2M (Sa – NaOH 2M) in both YPD and YPD with bromocresol purple (YPD + BP). Zoomed images from day 7 are displayed. White arrows indicate evident growth of new phenotypes within *S. aureus* original colonies. Sa – PBS represents the control condition. (B) Timeline of glucose effect on *S. aureus* variant colonies’ generation in a proximity assay of *S. aureus* with *C. albicans* (Ca – Sa) in both YPLD (YPD with glucose at 0.2 %) and YPLD with bromocresol purple (YPLD + BP). Sa – Sa is the control condition in which *S. aureus* inoculum was used in both columns, while Ca – Ca is the control condition where *C. albicans* inoculum was used in both columns. Displayed images are representative of three different experiments and were processed with ImageJ software. The scale bar of 2 cm applies to all cases. The shown distances correspond to P1= 1 cm, P2= 1.5 cm, P3= 2 cm, P4= 3 cm, P5= 4 cm, and P6= 5 cm.

Furthermore, we decided to test the role of glucose in the development of *S. aureus* colony variants, considering that glucose availability was reported to influence the Staphylococcal *agr* system (15) (Fig. 4B). A proximity assay between *S. aureus* and *C. albicans* in YPD with low glucose (YPLD) showed substantial variations in the growth of both microorganisms. Firstly, *C. albicans* growth decreased compared to YPD colonies, reflected in a smaller colony size at the end of the experiment in the control (Ca – Ca) and, more markedly, in the interaction with *S. aureus* (Ca – Sa). In contrast, *S. aureus* colonies growing in YPLD expanded overtime, reaching a colony size considerably higher than when they grew in YPD, regardless of the presence of *C. albicans*. Additionally, *S. aureus* growth was not accompanied by the acidification of the media, resulting in the total dominance of *C. albicans* alkalinization and the pH neutralization of the complete plate. These changes did not promote the generation of *S. aureus* colony variants.

### *S. aureus* colony variants exhibit increased virulence

To assess virulence differences of *S. aureus* colony variants, we conducted a proximity assay for 10 days. Each *S. aureus* colony was designated with a number, including controls (Fig. 5A). Colonies 1 to 4 corresponded to the new variants gradually generated at P1-P3 positions, where colony 4 represent the phenotype developed after initial exposure to basic pH, and colony 1 represented the phenotype obtained after sustained exposure. Colony 5 and 6 corresponded to the border and center of *S. aureus* growth at positions P4-P6, where no meaningful pH change was present. Controls were taken from a Sa – Sa plate incubated for 10 days (colony 7) and 1 day (colony 8).

**FIG 5.**
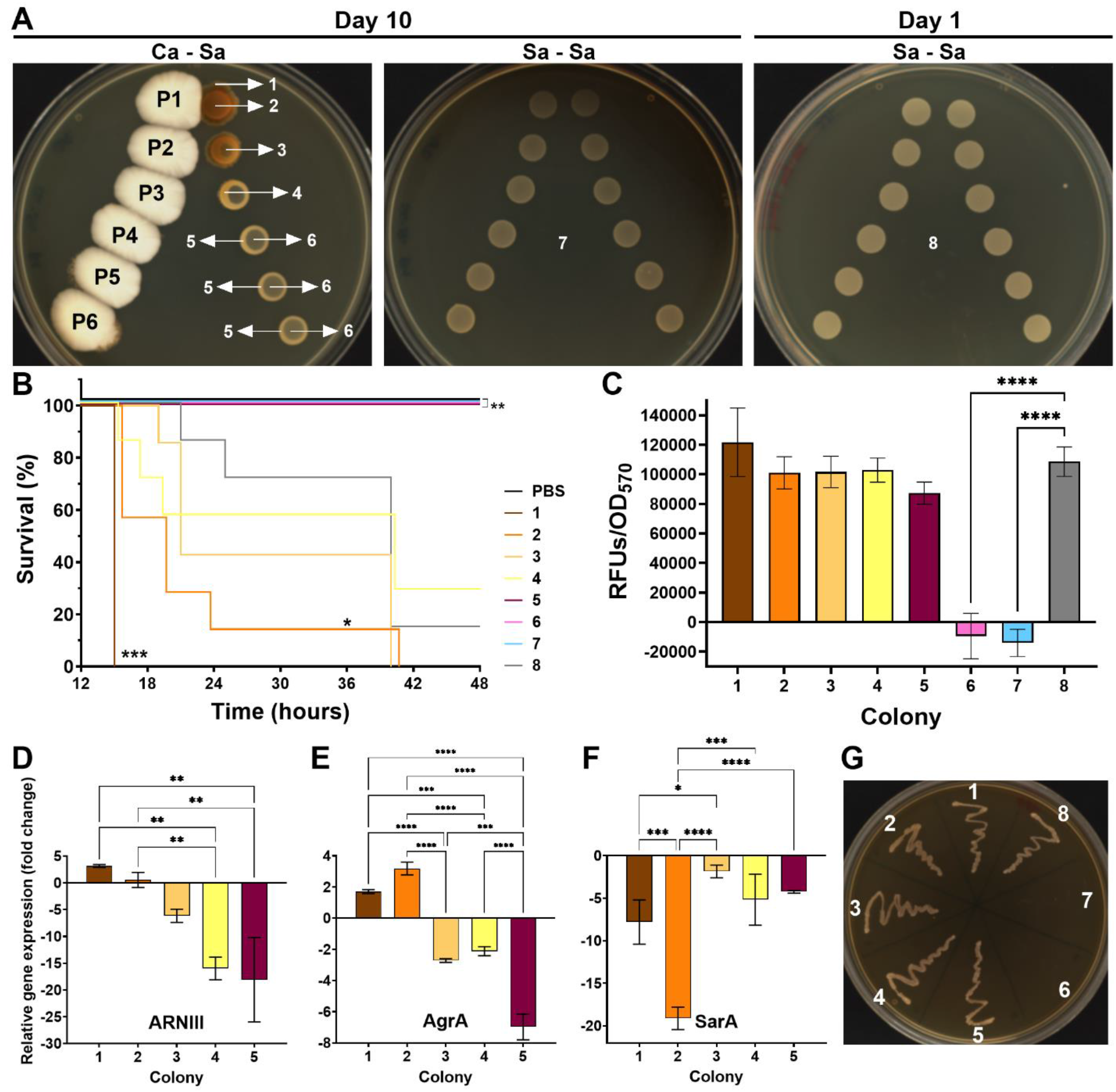
Virulence assessment of *S. aureus* colony variants. (A) Colonies of *S. aureus* obtained after a proximity assay with *C. albicans* (Ca – Sa) for 10 days were numbered. Colonies obtained from *S. aureus* controls (Sa – Sa) at day 10 and day 1 were also included. The shown distances correspond to P1= 1 cm, P2= 1.5 cm and P3= 2 cm, P4= 3 cm, P5= 4 cm, and P6= 5 cm. (B) Kaplan-Meier survival curves of *G. mellonella* larvae after inoculation with all different *S. aureus* colonies, with each condition involving 8 larvae. Asterisks indicate statistically significant differences in virulence in comparison to control colony 8 by a long-rank test. (C) Evaluation of the metabolic activity of *S. aureus* colonies by PrestoBlue assay. Asterisks indicate statistically significant differences versus control colony 8 in an Ordinary one-way ANOVA with Šidák’s multiple comparison test. RFUs= Relative Fluorescence Units. Fold change of (D) ARNIII, (E) AgrA, and (F) SarA gene expression of *S. aureus* colonies 1-5 in comparison to control colony 8. Asterisks indicate statistically significant differences among colonies in an Ordinary one-way ANOVA with Šidák’s multiple comparison test. (G) Reversion test of *S. aureus* colonies. The results presented in this figure are representative of the same experiment repeated three times. Images were processed with ImageJ software and error bars display mean and standard deviation. ∗*p*-value <0.05, ∗∗*p*-value <0.01, ∗∗∗*p*-value <0.001, and ∗∗∗∗*p*-value <0.0001

Infection of *G. mellonella* with *S. aureus* colonies showed significant differences in larval survival (Fig. 5B). Larvae infected with colonies 5, 6 and 7 did not show mortality. Meanwhile, the median survival of larvae infected with colonies 1, 2, 3, 4 and 8 was 15 h, 19 h, 21 h, 40 h and 40 h, respectively. According to the Log rank test, colonies 1 and 2 exhibited enhanced virulence compared to control colony 8. These variations cannot be attributed to differences in metabolic activity, as no statistically significant deviation was observed in the PrestoBlue results between colonies 1-5 and the control colony 8 (Fig. 5C). However, the complete survival of larva infected with colonies 6 and 7 was justified by their lack of metabolic activity.

Thus, the expression of three important virulence genes of *S. aureus* was evaluated in colonies 1-5 compared to the control colony 8 (Fig. 5D-F). We found that colony 1, the most virulent, exhibited increased expression of *ARNIII* and *agrA* but decreased expression of *sarA*. This induction of *agrA* and reduction of *sarA* expression were more pronounced in colony 2, although no change in RNAIII was observed in this case. Colonies 3-5 showed decreased expression of all genes; however, repression of the *agrA* gene was higher in colony 5 compared to colonies 3 and 4. Therefore, *S. aureus* variant colonies (colonies 1-4) originating in response to external pH changes exhibited increased virulence, associated to expression changes, compared to the original colonies (colonies 5-6) and the colonies formed under control conditions (Sa – Sa) for 10 days (colony 7) and 1 day (colony 8).

Finally, the re-plating of the colonies showed no differences among their growth (Fig. 5G), indicating that the colony variant phenotype is reversible.

## Discussion

The ability of both *C. albicans* and *S. aureus* to colonize a wide variety of human body sites increases the likelihood of shared host niches and the development of mixed infections (7). Survival in different microenvironments requires metabolic flexibility controlled by time- and space-specific regulatory networks (17, 18). This implies that environmental factors can influence disease outcomes (13). Moreover, in the context of polymicrobial infections, the environmental response of one microorganism can affect the behavior of other species (13).

In the case of *S. aureus*, the regulatory network controlling virulence responds to various environmental cues that drive the bacterium throughout the infection process (17). For instance, the *agr* system, responsible for ARNIII expression and α-toxin secretion, is sensitive to acidic or alkaline conditions, with a maximum activity near neutral pH (1, 14, 15). Furthermore, *S. aureus* virulence is enhanced by *C. albicans,* resulting in what is described as lethal synergism (9). Recent studies have shown that this effect is associated with an increase in *S. aureus* α-toxin production via *agr* activation, in response to extracellular ribose depletion and media neutralization caused by *C. albicans* (3, 12, 13). However, questions regarding this cooperative microbial interaction mediated by environmental crosstalk remain.

In this study, we performed a proximity assay that allowed us to monitor colony changes resulting from non-physical interactions between *S. aureus* and *C. albicans.* In contrast to previous studies that used liquid conditions for co-culture, our experiment permitted the simultaneous evaluation of six different distances between species growth. Long-term plate incubation (over 9-10 days) compensated for the higher diffusion times and loss of volatile compounds present in solid agar, allowing events to be recorded sequentially. As a result, our investigation reports that *C. albicans* enhances *S. aureus* virulence by inducing the generation of successive new colony variants (Fig. 1 and Fig. 5). The origin of the colony variants coincided with the pH change induced by *C. albicans*-mediated media alkalinization (Fig. 3), although *C. albicans* is not strictly required for their generation, as their appearance under other artificial pH modulators (NaOH) was also confirmed (Fig. 4).

Importantly, *G. mellonella* larvae survival (Fig. 5B) decreased in relation to the generation time of the colony variant. Colonies generated after prolonged exposure to *C. albicans* proximity (colonies 1-2) caused significantly higher mortality compared to those exposed for a shorter time (colonies 3-4) or not exposed at all (colonies 5 and 8) (Fig. 5A-B). This agrees with the lethal synergism observed from *in vivo* experiments of systemic and local infections after co-inoculation of both microorganisms (3, 9–11, 19, 20). Our results showed that *G. mellonella* larvae survival at 24 h decreased from 71.4% when infected with colony 8 to 0- 14% when infected with colonies 1 and 2, respectively. These values are consistent with the previous study by Sheehan *et al.* (2020) using this animal model, where larvae infected with *S. aureus* alone and in combination with *C. albicans* showed a survival of 80% vs 30% at 24 h (10).

Moreover, our experimental setting allowed us to recreate the interaction between both species, but to evaluate the virulence of *S. aureus* colonies by themselves. The isolation of the different *S. aureus* colonies from the proximity assay before larvae inoculation demonstrated that, although *C. albicans* is the responsible for the increase of *S. aureus* virulence, it is not required for the lethal effect. In addition, the qRT-PCR results showed that the increase in *S. aureus* virulence is related to an up-regulation of *ARNIII* and *agrA*, two important genes controlled by the *agr* system (Fig. 5D-E). Therefore, we believe the high pathogenicity of *S. aureus* colony variants originated after *C. albicans* exposure is related to an increase in α-toxin expression, as previously reported (3), although we did not perform a direct detection of this molecule.

Considering our recent discoveries and those of others, we propose a comprehensive model of *C. albicans*-*S. aureus* metabolic interactions and microenvironmental changes that could result in the enhancement of *S. aureus* virulence through new variants generation (Fig. 6). First, the aerobic growth of *C. albicans* under high glucose couples the glycolysis pathway with the TCA cycle (21). Although glucose is its preferred carbon source, this yeast can simultaneously metabolize alternative sources through enzymes from the gluconeogenic and glyoxylate cycle (22–25). Meanwhile, *S. aureus* aerobic growth in glucose rich-media causes repression of the tricarboxylic acid (TCA) cycle by CcpA. Thus, pyruvate obtained from glycolysis is converted to acetate and excreted into the media, causing pH decrease and downregulating the staphylococcal *agr* system (15, 18).

**Figure 6.**
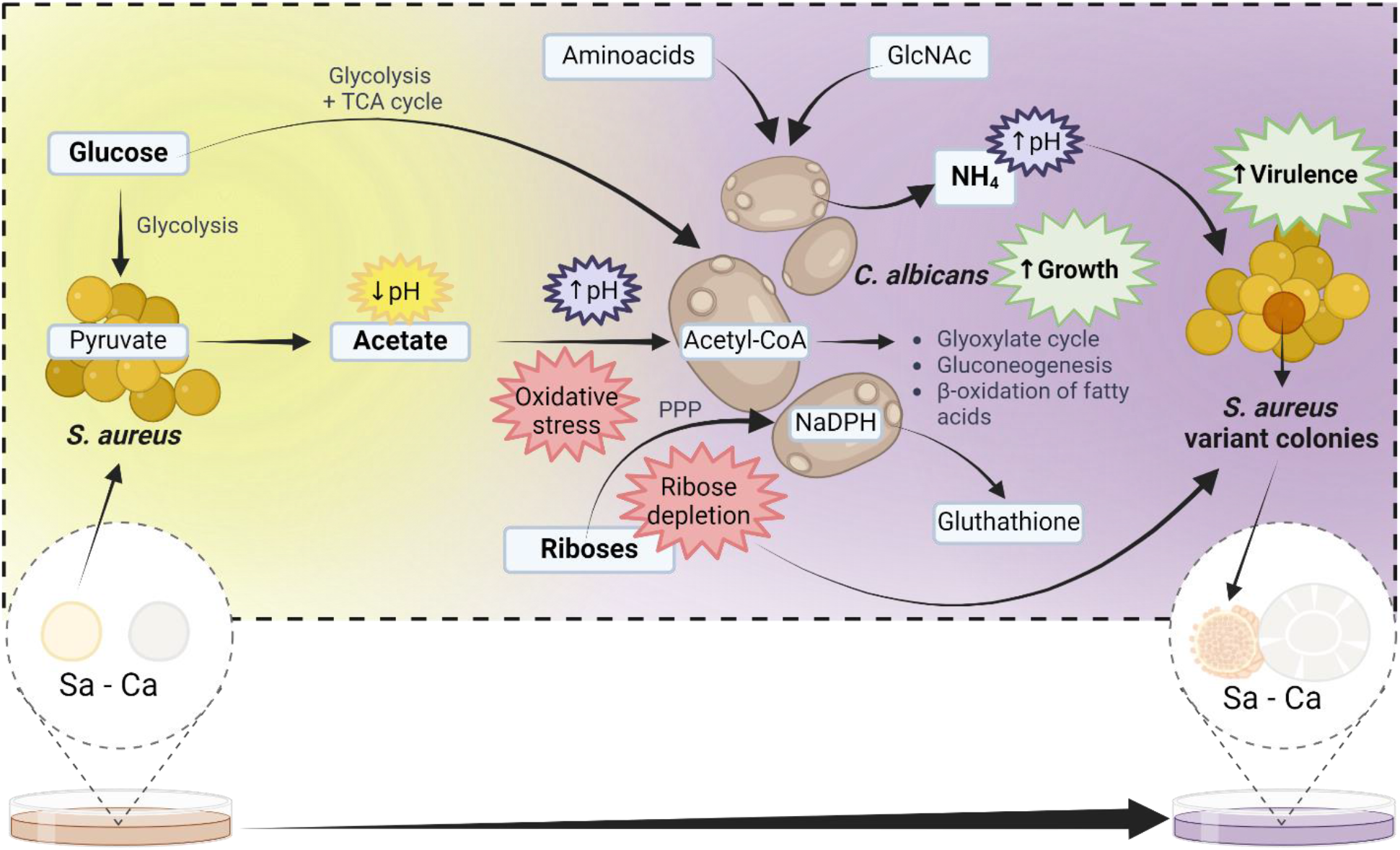
Working model of *S. aureus* and *C. albicans* metabolic interactions and their associated environmental changes influencing *S. aureus* virulence. *S. aureus* catabolism of glucose produces the excretion of acetate into the media, which decreases the pH. The acetate molecule serves as a precursor of Acetyl-coA, positively impacting the growth of *C. albicans*. Acetate consumption through different metabolic routes helps neutralize the pH of the media, in conjunction with ammonia (NH_4_) production derived from amino acids and/or N-acetylglucosamine (GlcNAc) metabolism. Additionally, extracellular acetate can induce oxidative stress in *C. albicans* cells, leading to ribose metabolism in the Pentose Phosphate Pathway (PPP) to produce the required NADPH for glutathione synthesis. pH neutralization and ribose depletion of the media by *C. albicans* activate the Staphylococcal *agr* system, increasing *S. aureus* virulence. TCA= Tricarboxylic Acids. Image created with Biorender.com.

Acetate may serve as source of acetyl-coA, the central intermediate in carbon metabolism (26). It is naturally present in blood and within macrophages and neutrophils (26). Once glucose is depleted, *C. albicans* can utilize acetate as a carbon source, entering the glyoxylate cycle, passing through the TCA cycle, and ultimately producing glucose in the gluconeogenic pathway (21). Additionally, acetate at physiological relevant concentrations up-regulates enzymes related to fatty acid metabolism (26). Therefore, this could be an explanation of the *S. aureus* positive impact on *C. albicans* growth (as observed in Fig. S1).

Furthermore, we believe that the consumption of acetate by *C. albicans* has an alkalinizing effect on the media, as reported with the metabolism of other carboxylic acids (α- ketoglutarate, pyruvate, and lactate) (27). However, the environmental changes in pH exerted by *C. albicans* may also involve two alternative mechanisms that this yeast has evolved to counteract acidic environments (27). The capacity to manipulate extracellular pH is related to survival inside phagocytic cells of the immune system and hyphal morphogenesis, being essential for fungal fitness *in vitro* and *in vivo* (16, 27). Both metabolism of amino acids and N-acetylglucosamine results in ammonia externalization, leading to a pH increase in the media (7, 16, 28, 29). This pH neutralization offers optimal conditions for the staphylococcal *agr* system, inducing α-toxin production (7).

Recently, the work of the field experts Peters and colleagues discovered a fundamental role of *C. albicans* ribose catabolism in activating the *agr* system of *S. aureus* (12). Although our experiments do not evaluate this new factor, as were performed before the publication of this finding, we propose that *C. albicans* ribose catabolism may be related to the production of NADPH by the Pentose Phosphate Pathway as a response to counteract the oxidative damage induced by acetate. Physiological relevant concentrations of acetate have been shown to induce oxidative stress in *C. albicans* cells, leading to glutathione secretion into the media, and NADPH is an important molecule in the glutathione redox cycle (26, 30, 31).

Bringing all these points together suggests that the lethal synergism observed in *C. albicans* and *S. aureus* co-infection results from a series of metabolic interactions that include environmental changes to which both species react. This is supported by the phenotype reversion of the colony variants after replating (Fig. 5G) once the corresponding stimuli were removed. However, we do not yet understand the evolutionary advantage that allows *S. aureus* cells that progressively adapt the transcription of virulence genes (*agr* system) in response to environmental stimuli (pH and ribose depletion) to have greater fitness, generating the appearance of colony variants. Nevertheless, evidence of *S. aureus* interconnection between metabolism and virulence is increasing daily, with more metabolic transcriptional regulators influencing the expression of virulence factors described each time (18).

Finally, we highlight the utility of the proximity assay as a simple, inexpensive, and easy-to-implement technique for studying visible colony changes resulting from non-physical interactions among microorganisms. Moreover, its use as a screening technique allowed us to identify similar interactions between *C. albicans* and two other bacterial pathogens, *S. epidermidis* and *E. coli*, suggesting potential avenues for future experiments. Additionally, we found that the *C. parapsilosis* clinical isolate used in this study can alkalinize the pH (data not shown) and therefore induce the generation of *S. aureus* colony variants.

## Materials and methods

### Bacterial strains and growing conditions

The fungaemia clinical isolates *Candida albicans* 10045727 and *Candida parapsilosis* 11103595 (32), and the bacterial strains *Staphylococcus aureus* ATCC 12600 (CECT 86), *Staphylococcus epidermidis* ATCC 1798 (CECT 231), *Streptococcus mutans* ATCC 25175 (CECT 479), *Enterococcus faecalis* ATCC 19433 (CECT 481) and *Escherichia coli* CFT073 (ATCC 700928) were used in this study. Each microorganism was recovered from a -80 stock. Bacteria were cultured at 37 °C in Tryptic Soy Agar (TSA) (Scharlab S.L.) or Luria Bertani (LB) agar (Scharlab S.L.), while fungi were grown at 30 °C in Yeast Petone Dextrose (YPD) medium consisting of 1% Yeast Extract (Gibco), 2% Meat Peptone (Scharlab S.L.), 2% D-glucose (Fisher Scientific S.L.), and, when required, 2% Bacteriological Agar (Scharlab S.L.). Similarly, overnight cultures were obtained after 16 h of incubation at 200 rpm and 30 °C in YPD for *Candida* and 37 °C in Tryptic Soy Broth (TSB) (Scharlab S.L.) or LB broth for bacterial strains.

### Proximity assays

Overnight cultures of *C. albicans* and *S. aureus* were centrifuged at 4000 rpm (Labnet Spectrafuge™ 6C) for 5 min and washed twice with Phosphate Buffer Saline 1X (PBS) (Fisher Scientific S.L.). Suspensions with a final optical density of λ = 550 nm (OD_550_) of 0.1 for yeast and 0.005 for bacteria (∼1x10^6^ CFUs/mL of both microorganisms) were prepared in PBS. Then, a 5 µl drop of each suspension was deposited in two parallel columns on YPD agar in a manner where the distance between both microorganisms drops increased progressively (Ca – Sa). In total, six different distances were tested: P1= 1 cm, P2= 1.5 cm, P3= 2 cm, P4= 3 cm, P5= 4 cm, and P6= 5 cm. Petri dishes were incubated at 37 °C for 9 days, and colony growth were recorded by imaging with the colony counter ProtoCOL 3 (Synbiosis). All images were processed with ImageJ software. Controls were created by facing drops from the same microorganism (Sa – Sa and Ca – Ca). Experiments were conducted thrice with 3 technical replicates each time. The same procedure was followed using YPD agar with Bromocresol purple at 0.08 mg/ml, denoted as YPD + BP, and YPD agar prepared with low D-glucose content (0.2 %), referred as YPLD.

Finally, we used a slightly modified proximity assay to confirm the pH influence on *S. aureus* colonies. First, a 5 µl drop from an *S. aureus* suspension was deposited on the agar in each corresponding position, leaving the facing positions empty. From day 1 to day 3, a 10 µl drop of a solution of NaOH 2M was applied in the morning and afternoon to the facing positions. On day 4, the quantity of the drop was increased to 15 µl, and on day 6, to 20 µl. *S. aureus* responses were monitored until day 7. Control plates followed the same procedure but with addition of PBS.

To confirm the specificity of the observed interactions between *C. albicans* and *S. aureus*, additional proximity assays were conducted using both microorganisms against different species (*S. epidermidis*, *S. mutans*, *E. coli*, *E. faecalis* and *C. parapsilosis*). Required controls were also included.

### Virulence assessment of *S. aureus* colony variants

Differences in the virulence of the obtained colonies of *S. aureus* growing in the different positions of the proximity assay with *C. albicans* were evaluated using the animal model *G. mellonella.* Maintenance and injection of *G. mellonella* larvae were carried out as previously reported by our laboratory (33).

Six different colonies of *S. aureus* from the proximity assays with *C. albicans* (Ca – Sa) were obtained after 10 days of incubation. Control colonies were obtained from control plates of *S. aureus* – *S. aureus* (Sa – Sa) after 10 days and 1 day of incubation. Cells were recovered with a pipette tip, resuspended in PBS until an OD_550_= 1.8, and injected into the hemocoel of eight larvae using a 26-gauge microsyringe (Hamilton). PBS-treated larvae were included as control. The experiment was conducted three times, and larvae were monitored for a total of 48 h post-injection with intervals of 2h from 15 h-25 h and 40-48 h.

At the same time, 100 µl of *S. aureus* cells suspensions were centrifugated (4000 rpm x 1 min) and resuspended in a solution 1:10 of the resazurin-based reagent PrestoBlue in TSB media. After 30 min at 37°C in the dark, the metabolic activity of the colonies was measured by fluorescence (λExc= 535 nm and λem=635 nm) and OD_570_ in a SPARK Multimode microplate reader. The remaining volume of cells suspensions was mixed with 500 mL of the stabilization solution RNAlater (Invitrogen) and frozen at -20 °C until further processing.

### Characterization of gene expression of *S. aureus* colony variants

Cells suspensions of *S. aureus* colonies 1-5 obtained in a proximity assay with *C. albicans* (Ca – Sa) were used for RNA extraction using the GeneJET RNA Purfication Kit (Thermo Fisher Scientific). RNA obtained was treated with 10x TURBO DNase (Life technologies). DNA contamination was discarded by PCR amplification of the 16S rRNA housekeeping gene. RNA quantification was performed on a NanoDrop 1000 Spectrophotometer (Thermo Fisher Scientific). Reverse transcription of RNA was performed by mixing the Maxima Reverse Transcriptase (Thermo Fisher Scientific) with Random Hexamer primers (Thermo Fisher Scientific) according to the manufacturer’s instructions. The obtained cDNA was stored at -20 °C until use.

Quantitative real-time PCR (qRT-PCR) was performed with the PowerUp SYBR Green Master Mix (Applied Biosystems) in a StepOnePlus™ Real-Time PCR System (Applied Biosystems) according to the manufacturer’s protocol. The sequences of primers used for gene amplification can be consulted at Vaudaux et al., 2002 (34). Relative expression of *RNAIII*, *agrA* and *sarA* genes were calculated using the values from control colony 8.

### Reversion test of *S. aureus* colony variants

A small inoculum of each colony of *S. aureus* were taken at the end of a proximity assay (Ca – Sa) with a pipette tip and plated on a new YPD plate. Same procedure was performed with control colonies (Sa – Sa). After 24 h of incubation, the growth of the colonies was evaluated.

### Statistical analysis

All data were statistically analyzed using GraphPad Prism version 10.00 (GraphPad Software, USA). Comparison of means among groups was performed by One-way ANOVA tests with a Šídák’s multiple comparisons test. Comparison of Kaplan-Meier survival curves were made by Log-rank tests. A *p*-value <0.05 was considered statistically significant.

## Acknowledgments

We thank Dr Jesus Guinea Ortega from the Department of Clinical Microbiology and Infectious Diseases, Hospital General Universitario Gregorio Marañón, Madrid, Spain, for the generous gift of the *Candida albicans* 10045727 and *Candida parapsilosis* 11103595 isolates used in this study.

This study was partially supported by grants PID2021-125801OB-100, PLEC2022- 009356 and PDC2022-133577-I00 funded by MCIN/AEI/ 10.13039/501100011033 and “ERDF A way of making Europe”, the CERCA programme and *AGAUR-Generalitat de Catalunya* (2021SGR01545), the European Regional Development Fund (FEDER) and Catalan Cystic Fibrosis association. The project that gave rise to these results received the support of a fellowship from ”la Caixa” Foundation (ID 100010434). The fellowship code is “LCF/BQ/DI20/11780040”.

Funding sources were not involved in the research conduction or article preparation.

## Declarations

**Conflict of Interest** All authors declare that they have no conflict of interest.

**Data availability** The datasets generated during and/or analysed during the current study are available from the corresponding author on reasonable request.

## References

1. Todd OA, Peters BM. 2019. *Candida albicans* and *Staphylococcus aureus* pathogenicity and polymicrobial interactions: Lessons beyond Koch’s postulates. J Fungi 5:81.

2. Carolus H, Van Dyck K, Van Dijck P. 2019. *Candida albicans* and *Staphylococcus* Species: A threatening twosome. Front Microbiol 10:2162.

3. Todd OA, Fidel PL, Jr., Harro JM, Hilliard JJ, Tkaczyk C, Sellman BR, Noverr MC, Peters BM. 2019. *Candida albicans* augments *Staphylococcus aureus* virulence by engaging the Staphylococcal *agr* quorum sensing system. mBio 10:e00910–19.

4. Kong EF, Tsui C, Kucharíková S, Andes D, Van Dijck P, Jabra-Rizk MA. 2016. Commensal protection of *Staphylococcus aureus* against antimicrobials by *Candida albicans* biofilm matrix. mBio 7:e01365–16.

5. Kean R, Rajendran R, Haggarty J, Townsend EM, Short B, Burgess KE, Lang S, Millington O, Mackay WG, Williams C, Ramage G. 2017. *Candida albicans* mycofilms support *Staphylococcus aureus* colonization and enhances miconazole resistance in dual-species interactions. Front Microbiol 8:244638.

6. Lin YJ, Alsad L, Vogel F, Koppar S, Nevarez L, Auguste F, Seymour J, Syed A, Christoph K, Loomis JS. 2013. Interactions between *Candida albicans* and *Staphylococcus aureus* within mixed species biofilms. BIOS 84:30–39.

7. Eichelberger KR, Cassat JE. 2021. Metabolic adaptations during *Staphylococcus aureus* and *Candida albicans* co-infection. Front Immunol 12:797550.

8. Peters BM, Jabra-Rizk MA, O’May GA, Costerton JW, Shirtliff ME. 2012. Polymicrobial interactions: impact on pathogenesis and human disease. Clin Microbiol Rev 25:193–213.

9. Carlson E. 1982. Synergistic effect of *Candida albicans* and *Staphylococcus aureus* on mouse mortality. Infect Immun 38:921–924.

10. Sheehan G, Tully L, Kavanagh KA. 2020. *Candida albicans* increases the pathogenicity of *Staphylococcus aureus* during polymicrobial infection of *Galleria mellonella* larvae. Microbiol 166:375–385.

11. Wu Y-M, Huang P-Y, Cheng Y-C, Lee C-H, Hsu M-C, Lu J-J, Wang S-H. 2021. Enhanced virulence of *Candida albicans* by *Staphylococcus aureus*: Evidence in clinical bloodstream infections and infected zebrafish embryos. J Fungi 7:1099.

12. Saikat P, Olivia AT, Kara RE, Christine T, Bret RS, Mairi CN, James EC, Paul L Fidel, Jr., Brian MP. 2024. A fungal metabolic regulator underlies infectious synergism during *Candida albicans*-*Staphyloccocus aureus* intra-abdominal co-infection. bioRxiv 2024.02:15.580531.

13. Todd OA, Noverr MC, Peters BM. 2019. *Candida albicans I*mpacts *Staphylococcus aureus* alpha-toxin production via extracellular alkalinization. mSphere 4:10.1128.

14. Regassa LB, Betley MJ. 1992. Alkaline pH decreases expression of the accessory gene regulator (*agr*) in *Staphylococcus aureus*. J Bacteriol 174:5095–5100.

15. Regassa LB, Novick RP, Betley MJ. 1992. Glucose and nonmaintained pH decrease expression of the accessory gene regulator (*agr*) in *Staphylococcus aureus*. Infect Immun 60:3381–3388.

16. Vylkova S, Lorenz MC. 2014. Modulation of phagosomal pH by *Candida albicans* promotes hyphal morphogenesis and requires Stp2p, a regulator of amino acid transport. PLOS Pathog 10:e1003995.

17. Jenul C, Horswill AR. 2019. Regulation of *Staphylococcus aureus* Virulence. Microbiol Spectr 7:10.1128.

18. Richardson AR. 2019. Virulence and Metabolism. Microbiol Spectr 7:10.1128.

19. Hu Y, Niu Y, Ye X, Zhu C, Tong T, Zhou Y, Zhou X, Cheng L, Ren B. 2021. *Staphylococcus aureus* synergized with *Candida albicans* to increase the pathogenesis and drug resistance in cutaneous abscess and peritonitis murine models. Pathogens 10:1036.

20. Vila T, Kong EF, Montelongo-Jauregui D, Van Dijck P, Shetty AC, McCracken C, Bruno VM, Jabra-Rizk MA. 2021. Therapeutic implications of *C. albicans*-*S. aureus* mixed biofilm in a murine subcutaneous catheter model of polymicrobial infection. Virulence 12:835–851.

21. Wijnants S, Vreys J, Van Dijck P. 2022. Interesting antifungal drug targets in the central metabolism of *Candida albicans*. Trends Pharmacol Sci 43:69–79.

22. Miramón P, Lorenz MC. 2017. A feast for *Candida*: Metabolic plasticity confers an edge for virulence. PLOS Pathog 13:e1006144.

23. Sandai D, Yin Z, Selway L, Stead D, Walker J, Leach Michelle D, Bohovych I, Ene Iuliana V, Kastora S, Budge S, Munro Carol A, Odds Frank C, Gow Neil AR, Brown Alistair JP. 2012. The Evolutionary rewiring of ubiquitination targets has reprogrammed the regulation of carbon assimilation in the pathogenic yeast *Candida albicans*. mBio 3:10.1128.

24. Lorenz MC. 2013. Carbon catabolite control in *Candida albicans*: New wrinkles in metabolism. mBio 4:10.1128.

25. Williams RB, Lorenz MC. 2020. Multiple alternative carbon pathways combine to promote *Candida albicans* stress resistance, immune interactions, and virulence. mBio 11:10.1128.

26. Bayot J, Martin C, Chevreux G, Camadro J-M, Auchère F. 2023. The adaptive response to alternative carbon sources in the pathogen *Candida albicans* involves a remodeling of thiol- and glutathione-dependent redox status. Biochem J 480:197–217.

27. Danhof Heather A, Vylkova S, Vesely Elisa M, Ford Amy E, Gonzalez-Garay M, Lorenz Michael C. 2016. Robust extracellular pH modulation by *Candida albicans* during growth in carboxylic acids. mBio 7:10.1128.

28. Naseem S, Min K, Spitzer D, Gardin J, Konopka JB. 2017. Regulation of Hyphal Growth and N-Acetylglucosamine Catabolism by Two Transcription Factors in *Candida albicans*. Genet 206:299.

29. Vesely EM, Williams RB, Konopka JB, Lorenz MC. 2017. N-Acetylglucosamine metabolism promotes survival of *Candida albicans* in the phagosome. mSphere 2:10.1128.

30. Komalapriya C, Kaloriti D, Tillmann AT, Yin Z, Herrero-de-Dios C, Jacobsen MD, Belmonte RC, Cameron G, Haynes K, Grebogi C, de Moura APS, Gow NAR, Thiel M, Quinn J, Brown AJP, Romano MC. 2015. Integrative model of oxidative stress adaptation in the fungal pathogen *Candida albicans*. PLOS ONE 10:e0137750.

31. Wang Y, Cao Y-Y, Jia X-M, Cao Y-B, Gao P-H, Fu X-P, Ying K, Chen W-S, Jiang Y-Y. 2006. Cap1p is involved in multiple pathways of oxidative stress response in *Candida albicans*. Free Radic Biol Med 40:1201–1209.

32. Marcos-Zambrano LJ, Escribano P, Bouza E, Guinea J. 2014. Production of biofilm by *Candida* and non-*Candida spp.* isolates causing fungemia: Comparison of biomass production and metabolic activity and development of cut-off points. Int J Med Microbiol 304:1192–1198.

33. Moya-Andérico L, Admella J, Torrents E. 2021. A clearing protocol for *Galleria mellonella* larvae: Visualization of internalized fluorescent nanoparticles. New Biotechnol 60:20–26.

34. Vaudaux P, Francois P, Bisognano C, Kelley William L, Lew Daniel P, Schrenzel J, Proctor Richard A, McNamara Peter J, Peters G, Von Eiff C. 2002. Increased expression of clumping factor and fibronectin-binding proteins by hemB mutants of *Staphylococcus aureus* expressing small colony variant phenotypes. Infect Immun 70:5428–5437.

